# Positive allosteric modulation of a GPCR ternary complex

**DOI:** 10.1101/2024.04.04.588200

**Authors:** Wessel A.C. Burger, Christopher J. Draper-Joyce, Celine Valant, Arthur Christopoulos, David M. Thal

**Author notes:** **Corresponding Authors** Celine Valant, Arthur Christopoulos, David M. Thal.

## Abstract

The activation of a G protein-coupled receptor (GPCR) leads to the formation of a ternary complex between agonist, receptor, and G protein that is characterised by high-affinity binding. Allosteric modulators bind to a distinct binding site from the orthosteric agonist and can modulate both the affinity and the efficacy of orthosteric agonists. The influence allosteric modulators have on the high-affinity active state of the GPCR-G protein ternary complex is unknown due to limitations on attempting to characterize this interaction in recombinant whole cell or membrane-based assays. Here, we use purified M_2_ muscarinic acetylcholine receptor (mAChR) reconstituted into nanodiscs to show that once the agonist-bound high-affinity state is promoted by the G protein, positive allosteric modulators stabilise the ternary complex that, in the presence of nucleotides leads to an enhanced initial rate of signalling. Our results enhance our understanding of how allosteric modulators influence orthosteric ligand signalling and will aid the design of allosteric therapeutics.

**Teaser:** Allostery from top and bottom, the combined influence of positive allosteric modulators on receptor signalling.

## Introduction

G protein-coupled receptors (GPCRs) are cell surface proteins activated by extracellular environmental stimuli and, in response, facilitate the modulation of intracellular signalling pathways and biological responses (*1*). GPCRs are important drug targets, and understanding how different ligands and heterotrimeric G proteins interact and modulate GPCR activity can facilitate the design of future therapeutics (*2*). A milestone in GPCR pharmacology was the observation that agonists have low- and high-affinity binding states for a receptor that was dependent on the presence of guanine nucleotides (*3*). The low-affinity state (K_Low_) arises from the binding between an agonist and receptor, whereas the high-affinity state (K_High_) represents a ternary complex between agonist, receptor, and G protein, with the increase in binding affinity being due to positive cooperativity between the agonist and G protein (*4, 5*). These observations and others (*6*) form the basis of the ternary complex model (TCM) (*7*), a central paradigm in modern GPCR pharmacology that posits transducer proteins such as G proteins are endogenous positive allosteric modulators (PAMs) of agonist-GPCR complexes.

Structural biology studies have provided mechanistic insight into the interactions of agonists and G proteins with receptors (*8*–*10*). The binding of agonists to GPCRs promotes conformational rearrangements within GPCRs that activate the receptor and accommodate the intracellular binding of heterotrimeric G proteins and other transducer proteins (*10*). NMR experiments revealed that agonist binding alone does not fully stabilise the active receptor conformation and requires interaction with the G protein (*11*–*13*) supporting the TCM. Further interrogation of high-affinity agonist binding was made possible by reconstituting purified GPCRs into nanodiscs (*14, 15*). Nanodiscs provide a stable platform to interrogate the relative ratios of a GPCR to a G protein in a reductionist approach that allows the determination of the low-affinity (*K*_Low_) and high-affinity (*K*_High_) states away from the cellular milieu. Consistent with the TCM was the observation that shifts in agonist affinity (*K*_Low_/*K*_High_) promoted by transducers correlated strongly with molecular efficacy, suggesting that the transducer cooperativity can be used as a surrogate measure of efficacy (*16*). Pharmacological experiments using β_2_AR-nanodiscs showed that G proteins could alter the association and dissociation rates of orthosteric ligands, which implies an allosteric coupling between agonist and transducer binding, further validating the TCM (*17*).

Most GPCRs possess secondary, spatially distinct, binding sites known as allosteric sites (*9*). In recent years, there has been increased interest in the discovery and development of small molecule allosteric modulators that target allosteric sites because these binding sites offer the potential for the design of more selective ligands in comparison to orthosteric ligands that target conserved orthosteric sites (*18*–*20*). An exemplar system to study GPCR allostery is the muscarinic acetylcholine receptors (M_1_–M_5_ mAChRs) (*21*–*23*). The first PAM-bound structure was of the M_2_ mAChR in complex with a G protein-mimetic, the agonist iperoxo, and the PAM LY2119620 (LY211, Fig. 1A). The structure revealed that LY211 bound in a large allosteric binding site located above the orthosteric site in a solvent accessible extracellular vestibule (ECV) that was in a closed conformation (*24*). In another study, bitopic probes possessing an orthosteric agonist component and a component spatially restricted to the ECV, showed that the conformation of the ECV influenced the signalling capacity of the M_2_ mAChR. Similarly, a recent mutagenesis study showed that ACh affinity was enhanced when residues in the ECV were mutated (*25, 26*). Together, these studies and others highlight the allosteric coupling between allosteric and orthosteric binding sites.

**Figure 1:**
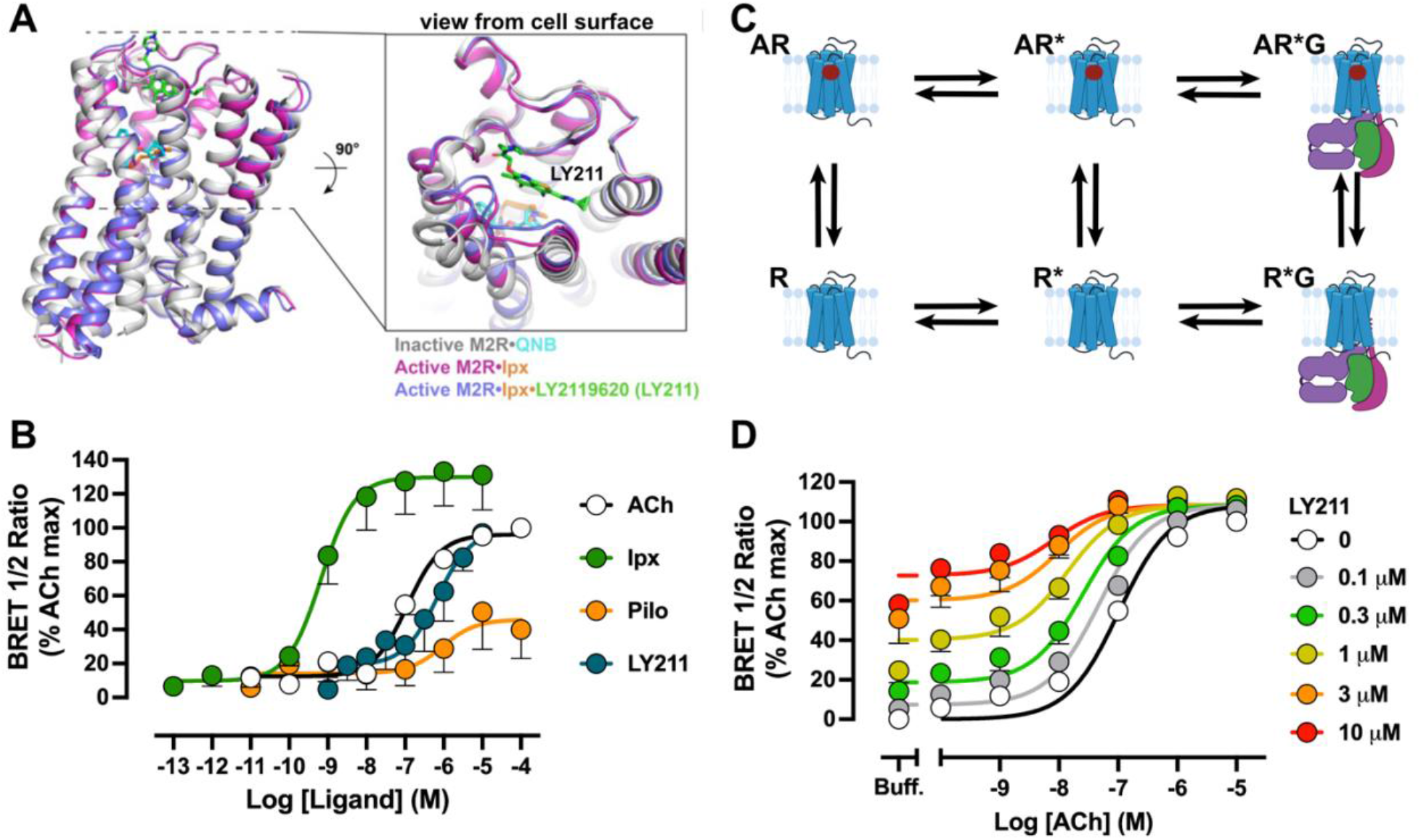
Allosteric receptor and G protein activation at the M_2_ mAChR. **(A)** Comparison of inactive M_2_ mAChR structure bound to QNB (PDB: 3UON, receptor-coloured white and QNB cyan), to active M_2_ mAChR structure bound to iperoxo (PDB: 4MQS, receptor-coloured magenta with Ipx in orange) and to active + PAM M_2_ mAChR structure bound to iperoxo and LY211 (PDB: 4MQT, receptor coloured blue with Ipx in orange and LY211 in green). **(B)** G_i1_ activation in M_2_ mAChR FlpIn CHO cells. Concentration-response curves of ACh, Ipx, Pilo and LY211. Data represent ligand induced-BRET normalised to the maximum response of Ipx fitted globally to a three-parameter logistic equation. **(C)** The extended ternary complex where A is agonist, G is G protein, R is an inactive receptor, and R* is the active receptor. **(D)** G_i1_ activation in M_2_ mAChR FlpIn CHO cells. Interaction of LY211 and ACh. Data were the ligand induced-BRET normalised to the maximum response of ACh, and were fitted globally to an operational model of allosterism. Data represent the mean ± S.E.M. of at least three individual experiments performed in duplicate. Group sizes and obtained parameters are listed in Table 1.

The combined effect of G proteins (endogenous PAMs) and exogenous small molecule PAMs on orthosteric agonist affinity and how this influences signalling is poorly established. In this study, we investigated how the PAM LY211 and G proteins influence agonist binding and signalling at GPCRs by reconstituting purified human M_2_ mAChRs into nanodiscs. The degree of cooperativity from the low-affinity to the high-affinity state promoted by the G protein was dependent on the efficacy of the orthosteric agonist and was not increased further by the addition of a PAM. This indicates that once the high-affinity state is promoted by the G protein, the PAM cannot further increase orthosteric agonist affinity. Investigation of the kinetics of the allosteric interaction between a PAM and the ternary complex revealed a decrease in the association and dissociation rates of the orthosteric ligand from the ternary complex in the presence of the PAM. By measuring the activation kinetics of G protein in the presence of allosteric modulators, we show that the PAM more readily stabilises the ternary complex, leading to an enhanced rate of signalling by the G protein. These results further our understanding of how small molecule PAMs affect the high-affinity state and GPCR signalling.

**Table 1.**
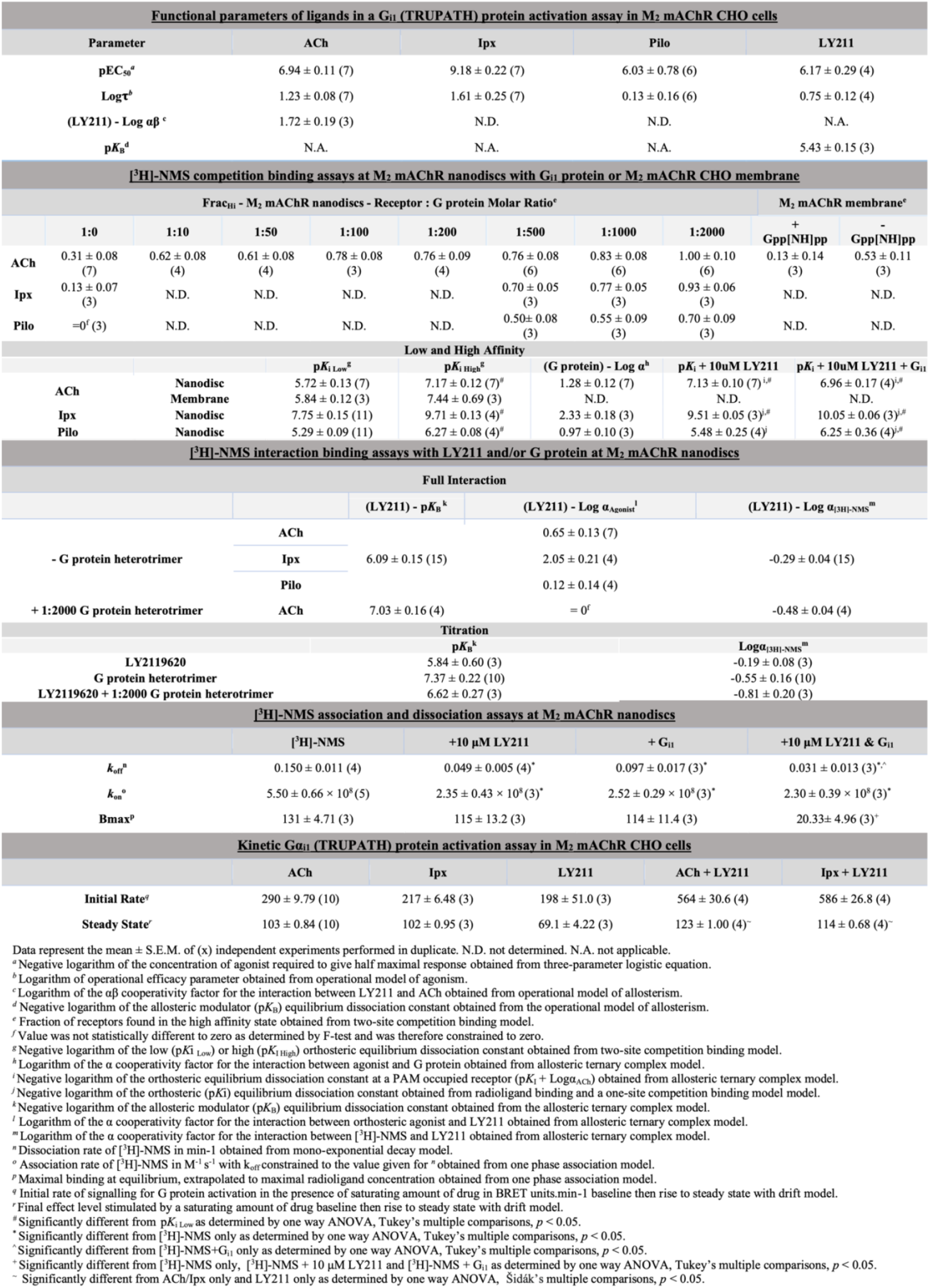
Pharmacological parameters from functional and radioligand binding experiments.

## Results

### G protein activation at the M_2_ mAChR

Following agonist binding, the agonist-bound receptor binds to and activates a G protein heterotrimer, leading to the dissociation of the Gα and Gβγ subunits (*27*). Wild type (WT) M_2_ mAChR expressing FlpIn Chinese Hamster Ovary (CHO) cells were transiently transfected with the TRUPATH bioluminescence resonance energy transfer (BRET) Gα_i1_β_3_γ_9_ sensors (*28*), and the decrease in BRET signal representative of subunit dissociation (and therefore G protein activation) in response to different agonists was measured. The orthosteric agonists ACh, iperoxo (Ipx), pilocarpine (Pilo), and the PAM-agonist (PAM-ago) LY211 promoted different extents of receptor-mediated G protein activation and efficacy (Fig. 1B, Table 1). As per the extended ternary complex model (ETCM), which builds on the TCM to account for receptor isomerization from the inactive to the active state (R to R*) prior to engagement of the R* state with G protein (R*G) (*29*) (Fig. 1C), this suggests that orthosteric and allosteric agonists differ in their ability to promote the conversion of AR to AR*. We considered ACh as the reference full agonist because it is the endogenous agonist. Ipx produced a maximal response greater than that of ACh and displayed greater efficacy (Table 1). For these reasons, Ipx was considered a ‘super agonist’ in this system, consistent with prior studies at the M_2_ mAChR (*30, 31*), indicating that Ipx has a greater capacity to form the AR* state than ACh. Conversely, Pilo is a partial agonist, as evidenced by its decreased efficacy relative to ACh (Table 1), and weakly promotes the conversion of the AR to AR* state. The PAM-agonist LY211 had a similar maximal response as ACh, illustrating that receptor activation can occur due to agonist binding at either the orthosteric or an allosteric site. Additionally, a concentration-response curve of ACh with increasing concentrations of LY211 revealed that ACh signalling was potentiated in the presence of LY211 (Fig. 1D). The data were analysed with an operational model of allosterism to derive a functional cooperativity factor (αβ) (*32*). The positive log αβ value indicates that LY211 enhances the ability of ACh to promote receptor-mediated G protein activation (conversion from R state to R* state) (Fig. 1D, Table 1). It is difficult to characterise the R*G state in functional assays due to the transient nature of this state in the presence of nucleotides; as such, we measured high-affinity binding (occurring at the R*G state) in radioligand binding experiments using M_2_ mAChR reconstituted into nanodiscs with purified WT G protein heterotrimer (Gα_i1_β_1_γ_2_).

### Reconstitution of the M_2_ mAChR into nanodiscs

To generate M_2_ mAChR nanodiscs, we used a modified receptor construct (M_2ΔICL3_ mAChR) that was also used in the x-ray crystallography and cryo-electron microscopy studies of the M_2_ mAChR. Expression and purification of M_2ΔICL3_ mAChR were performed as previously described (*24, 33*) (Fig. S1A-C). To verify that this receptor construct was functional, we stably transfected the M_2ΔICL3_ mAChR construct into CHO cells. Radioligand binding data indicated the subsequent cell line was low-expressing. Instead, we turned to a highly amplified functional assay and determined extracellular signal-regulated kinase 1/2 phosphorylation (pERK1/2) in response to ACh and Ipx (Fig. S1D). As expected for a low-expressing cell line, the obtained pEC_50_ values for ACh and Ipx resemble reported affinity values at the M_2_ mAChR. Furthermore, the rank order of agonism and potency, as well as the fold difference in potency between ACh and Ipx, was maintained between WT and M_2ΔICL3_ mAChR, validating the functionality of this receptor construct (*24*). Purified M_2ΔICL3_ mAChR was reconstituted into nanodiscs using a covalently circularized membrane scaffold protein (MSP), cMSP1D1 (*34*) (Fig. S2A-B) and the monomeric nature of the nanodisc was verified through complexing with an anti-Flag Fab and negative stain electron microscopy (Fig. S2C-E). Hereafter, we refer to reconstituted M_2ΔICL3_ mAChR in nanodiscs as M_2_ mAChR nanodiscs.

### Formation and characterisation of the high-affinity state at the M_2_ mAChR

To characterise the high-affinity state of M_2_ mAChR nanodiscs, we first expressed and purified a range of heterotrimeric G proteins with differing Gα subunits and Gβ_1_γ_2_ (Fig. S3A-B). At a 1:2000 molar ratio of M_2_ mAChR:G protein (R:G), G_i1_, G_i2_, G_i3_, G_oA_, and G_s_ heterotrimers were able to promote a similar increase in the affinity of ACh in radioligand competition binding experiments with [^3^H]-N-methylscopolamine (NMS) (Fig. S3C). G_q_ was not able to mediate any change in ACh affinity (Fig S3C). These results are consistent with our recent findings of G protein signalling at the M_2_ mAChR using the TRUPATH assay (*35*) and a recent study exploring the selectivity of nucleotide-free G proteins (*36*).

Given the lack of difference between canonical G_i/o_ proteins in increasing ACh affinity, we performed a complete characterisation of the high-affinity state with one of these G proteins, specifically G_i1_. Titrating amounts of G_i1_ heterotrimer at M_2_ mAChR nanodiscs increased the fraction of receptors in the high-affinity state (Frac_Hi_ values; Fig. 2A). At the 1:2000 R:G ratio, all of the receptor population was in the high-affinity state indicating this R:G ratio was sufficient for complete formation of the high-affinity state (Fig. 2E). Formation of the active state was accompanied by a loss of [^3^H]-NMS binding, consistent with destabilisation of the inactive conformation of the receptor. Analysing the low- and high-affinity states of ACh with a two-site competition binding model with p*K*_I(Low)_ and p*K*_I(High)_ values shared across the R:G ratios, revealed a 27-fold increase in the affinity of ACh between p*K*_I(Low)_ and p*K*_I(High)_ (Table 1). To verify that an incubation period of 6 hours at room temperature was sufficient for nucleotide-free G protein to bind M_2_ mAChR nanodiscs and reach equilibrium, we determined p*K*_i(Low)_ and p*K*_i(High)_ values following a shorter incubation period. Similar p*K*_i(Low)_ and p*K*_i(High)_ values were obtained at 4 hours, indicating that equilibrium was already reached before 6 hours (Fig. S4A) (*37*). To ensure receptor stability, we measured [^3^H]-NMS binding and determined nanodisc integrity through size-exclusion chromatography (SEC) over a 24-hour period at room temperature (Fig. S4B-C). Both approaches validated nanodisc stability and integrity up to 6 hours with the 24-hour time point leading to a slight loss in [^3^H]-NMS binding.

**Figure 2:**
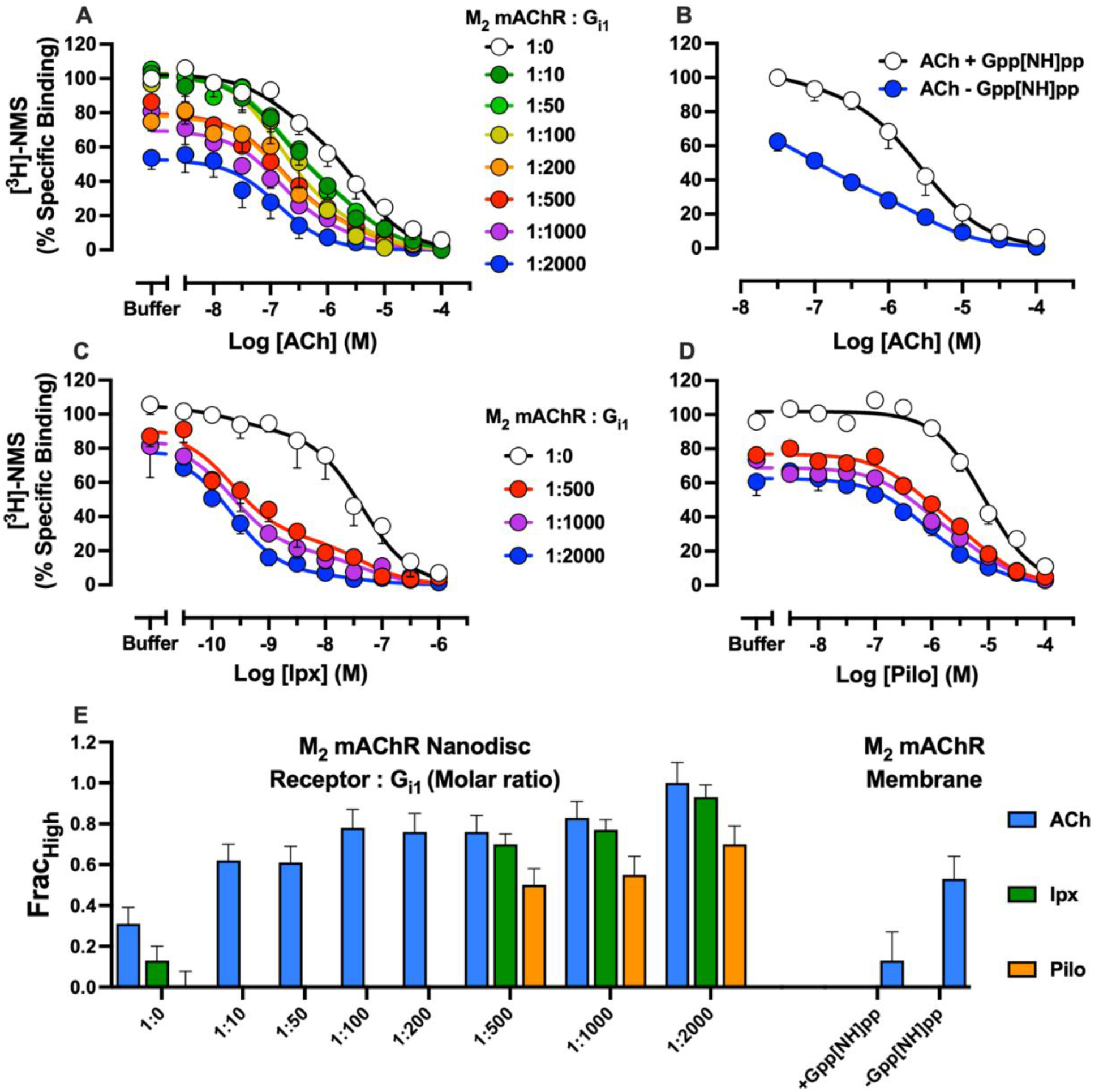
Modulation of the low and high-affinity state of orthosteric agonists by G protein. **(A)** Radioligand competition binding between ACh and [^3^H]-NMS at M_2_ mAChR nanodiscs in the presence of stoichiometric amounts of receptor to G_i1_ protein. **(B)** Radioligand competition binding between ACh and [^3^H]-NMS at M_2_ mAChR membranes with and without 100 μM Gpp[NH]pp. **(C)** Radioligand competition binding between Ipx and [^3^H]-NMS at M_2_ mAChR nanodiscs in the presence of stoichiometric amounts of receptor to G_i1_ protein. **(D)** Radioligand competition binding between Pilo and [^3^H]-NMS at M_2_ mAChR nanodiscs in the presence of stoichiometric amounts of receptor to G_i1_ protein. Data were normalised to the buffer only condition and were fitted globally to a two-state model of competition binding where p*K*_i (Low)_ and p*K*_i (High)_ values were shared. For all panels, data represent the mean ± S.E.M. of three individual experiments performed in duplicate. Group sizes and obtained parameters are listed in Table 1. **(E)** Fraction of M_2_ mAChR receptors found in the high-affinity state in the presence of different ligands, G protein ratios or Gpp[NH]pp from **A-D**.

To compare ACh affinity values at M_2_ mAChR nanodiscs to those observed in a more native cell environment, we performed competition binding between ACh and [^3^H]-NMS at M_2_ mAChR FlpIn CHO membranes (Fig. 2B). During membrane preparation, endogenous nucleotide is lost and must be added exogenously in the form of a GTP analogue, Gpp[NH]pp. By performing competition binding with and without Gpp[NH]pp, ACh affinity at receptor alone (p*K*_I (Low)_) and at the ternary complex (p*K*_I (High)_) can be determined (*4, 38*). In the absence of Gpp[NH]pp, around half of the receptors were in the high-affinity state (Fig. 2E), highlighting the difficulty in fully forming the high-affinity state in membranes compared to nanodiscs. Similar low- and high-affinity values were obtained with M_2_ mAChR membranes compared to M_2_ mAChR nanodiscs, justifying the use of nanodiscs to explore, in a more exclusive manner, the high-affinity state at the M_2_ mAChR (Table 1).

Given the difference in the ability of agonists to redistribute the R to R* equilibrium according to a G protein activation assay (Fig. 1B), we explored the impact of a super and partial agonist (Ipx and Pilo) on the high-affinity state. Competition binding between Ipx and Pilo vs [^3^H]-NMS were performed in the presence of the three highest stoichiometric amounts of G protein (Fig. 2C-E). For Ipx, similar Frac_Hi_ values as to those observed with ACh were seen (Fig. 2C,E), though there was a greater shift (91-fold) in Ipx affinity vs ACh (27-fold). In the presence of Pilo, a smaller fraction of receptors were in the high-affinity state compared to ACh, which is consistent with Pilo being a partial agonist (Fig. 2D,E). In contrast, there was a smaller shift (9.5-fold) from the low- to high-affinity state with Pilo (Table 1).

Considering that G proteins act as an endogenous allosteric modulator of agonist binding, we calculated binding cooperativity values (α) that describe the change in orthosteric ligand affinity when co-bound with an allosteric modulator using the allosteric ternary complex model (ACTM) (*19, 39*). Positive values indicate positive cooperativity, negative values indicate negative cooperativity and values of zero indicate neutral cooperativity. Consistent with our previously obtained p*K*_I(Low)_ – p*K*_I(High)_ values, G_i1_ displayed a binding cooperativity rank order of Pilo<ACh<Ipx (Table 1). Similarly, the collapse in radioligand binding observed with increasing G protein concentrations indicates the G protein acts as a NAM of antagonist binding. Plotting the G protein ratios (G_i1_) as a molar concentration and using the ATCM revealed a binding cooperativity value of log α of -0.55 ± 0.16 for [^3^H]-NMS binding and a binding affinity between the G protein and the NMS-bound receptor of approximately 42 nM (Fig. S6). Altogether, the differences between p*K*_I(Low),_ p*K*_I(High),_ Frac_Hi_, and (α) for agonists match the G protein activation assay results and validate that the degree of G protein activation and efficacy orthosteric agonists exhibit reflects the extent to which orthosteric agonists stabilise the active state. The difference in the direction of G protein cooperativity between agonists and antagonists reflects a two-state model of allostery and allows for the comparison of cooperativity between the endogenous G protein and exogenous allosteric modulators.

### G protein dictates the formation of the high-affinity state

Similar to G proteins, PAMs increase orthosteric ligand affinity. However, how this relates to the ability of the G protein to form the high-affinity state remains unknown. To explore this, we initially tested the ability of LY211 to modulate ACh, Ipx and Pilo affinity at M_2_ mAChR nanodiscs without the presence of G proteins in [^3^H]-NMS radioligand binding. Any increase in agonist affinity thus represents the ability of the PAM to modulate the low-affinity state of these agonists. Again, we used the ACTM (*19, 39*) to derive binding cooperativity values. LY211 is a PAM of ACh, Ipx and Pilo binding with a similar rank order compared to G protein, Pilo<ACh<Ipx (Fig. 3A, S5A-B, Table 1). As evidenced by the collapse in [^3^H]-NMS binding, LY211 negatively modulates [^3^H]-NMS binding (log α_[3H]-NMS_ = –0.29 ± 0.04). The observed cooperativity values for LY211 are consistent with previous studies (Table 1) (*24, 40*). In comparison to LY211, our data indicates that G_i1_ is a superior PAM of agonist binding while being a similar NAM of [^3^H]-NMS binding.

**Figure 3:**
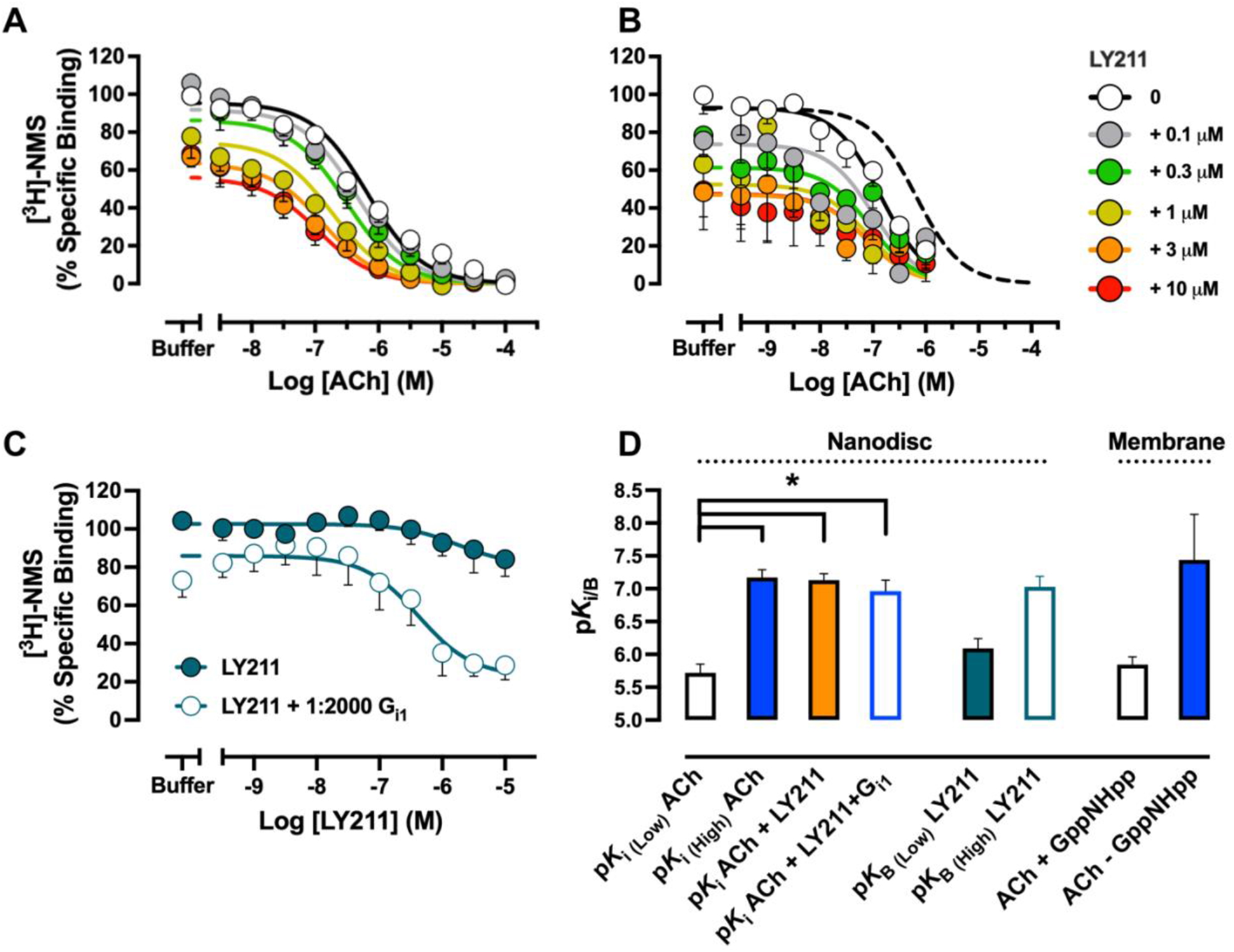
Modulation of the low and high-affinity state of orthosteric ligands by G protein and PAM. Radioligand competition binding between ACh and [^3^H]-NMS at M_2_ mAChR nanodiscs with increasing amounts of LY2119620 **(A)** without and **(B)** with a saturating 1:2000 stoichiometric R:G ratio of G_i1_ protein. The shift in ACh affinity in the presence of G_i1_ is illustrated through the inclusion of the ACh binding curve without G protein in **B** (dashed line). For both experiments, data were normalised to the buffer-only condition and were fitted globally to an allosteric ternary complex model. **(C)** Allosteric interaction between LY211 and [^3^H]-NMS at M_2_ mAChR nanodiscs with and without a saturating 1:2000 stoichiometric R:G ratio of G_i1_ protein. Data were normalised to the buffer-only condition and were fitted globally to an allosteric ternary complex model. **(D)** p*K*_i_ and p*K*_B_ values of ACh and LY211 were obtained in radioligand binding at M_2_ mAChR nanodiscs and M_2_ mAChR membranes. For all panels, data represent the mean ± S.E.M. of at least three individual experiments performed in duplicate. Group sizes and obtained parameters are listed in Table 1. *, significantly different, *p* < 0.05, one-way ANOVA, Tukey’s multiple comparisons test.

Next, we tested whether LY211 could modulate the high-affinity state and focused on ACh by performing an interaction between ACh and LY211 in the presence of a saturating amount of G protein, (R:G of 1:2000) (Fig. 3B). Increasing concentrations of LY211 produced no apparent change in the affinity of ACh, demonstrating that LY211 is a neutral allosteric ligand (NAL, log α_ACh_ = 0) of the high-affinity ACh-bound state. This further suggests that G_i1_ is a superior PAM compared to LY211, since no further modulation was observed. However, the addition of LY211 did promote a further collapse in [^3^H]-NMS binding, suggestive of an allosteric interaction between the G protein and the PAM for the antagonist-bound conformation. Indeed, given that LY211 and G_i1_ displayed similar cooperativity values for [^3^H]-NMS binding, it is not unreasonable to think the NAM effect could be additive. To explore more accurately, we titrated a wider range of LY211 concentrations against a K_D_ concentration of [^3^H]-NMS with and without a saturating amount of G_i1_ and determined the affinity (p*K*_B_) and log α_[3H]-NMS_ of LY211 through the use of the ATCM (Fig. 3C, Table 1). Without G protein, the affinity and cooperativity values resembled those obtained in the full interaction between [^3^H]-NMS, ACh, and LY211 as expected. In the presence of G protein, a 6-fold increase in the affinity of LY211 was observed as well as an increase in the ability of LY211 to allosterically displace [^3^H]-NMS (Fig. 3D, Table 1). Compared to the log α_[3H]-NMS_ of G protein only (Fig S6A, Table 1), this decrease when G protein and LY211 are combined indicates an additive NAM effect and additional destabilisation of the inactive state.

Because LY211 is a PAM-agonist that can promote the active state of the receptor on its own (*24, 40*) (Fig. 1B), we investigated what influence LY211 had on the high-affinity state of Ipx or Pilo. Particularly in the case of Pilo where low cooperativity values were observed for both G protein and PAM, we investigated whether the presence of both modulators would enhance Pilo’s ability to partially shift the AR to AR* receptor equilibrium. Since LY211 did not enhance ACh-bound high-affinity state (Fig. 3B), we measured the affinity of Ipx or Pilo in the presence of G protein without (p*K*_I (High))_ and with (p*K*_I (High+PAM_)) 10 μM LY211 only (open blue and closed blue circles, respectively (Fig.4 A-D). In this way, we could also perform 1) a competition binding curve of Ipx or Pilo without G protein (p*K*_I (Low),_ open black circles, Fig. 4A-D) and 2) Ipx and Pilo without G protein but with 10 μM LY211 (p*K*_I (Low+PAM),_ orange circles, Fig. 4A-D) in parallel. By performing the experiment in this manner, the influence that PAMs and G proteins have on agonist and [^3^H]-NMS affinity is more appropriately determined and compared. The presence of 10 μM LY211 significantly increased the affinity of Ipx by nearly 100-fold, similar to the influence of G protein on Ipx affinity (Fig. 4A,C, Table 1). The addition of both G protein and LY211 produced no significant increases in Ipx affinity beyond what either the G protein alone or PAM alone produced. In the case of Pilo, a non-significant increase of affinity was observed in the presence of 10 μM LY211. Similar to ACh and Ipx, the p*K*_I (High)_ of Pilo was not altered in the presence of PAM (Fig. 4B,D, Table 1). Thermodynamically, this indicates that the inherent efficacy of the orthosteric agonist in combination with the G protein, as a superior PAM, determines the formation of the high-affinity state and that small molecule PAMs cannot increase orthosteric ligand affinity once this high-affinity state is formed.

**Figure 4:**
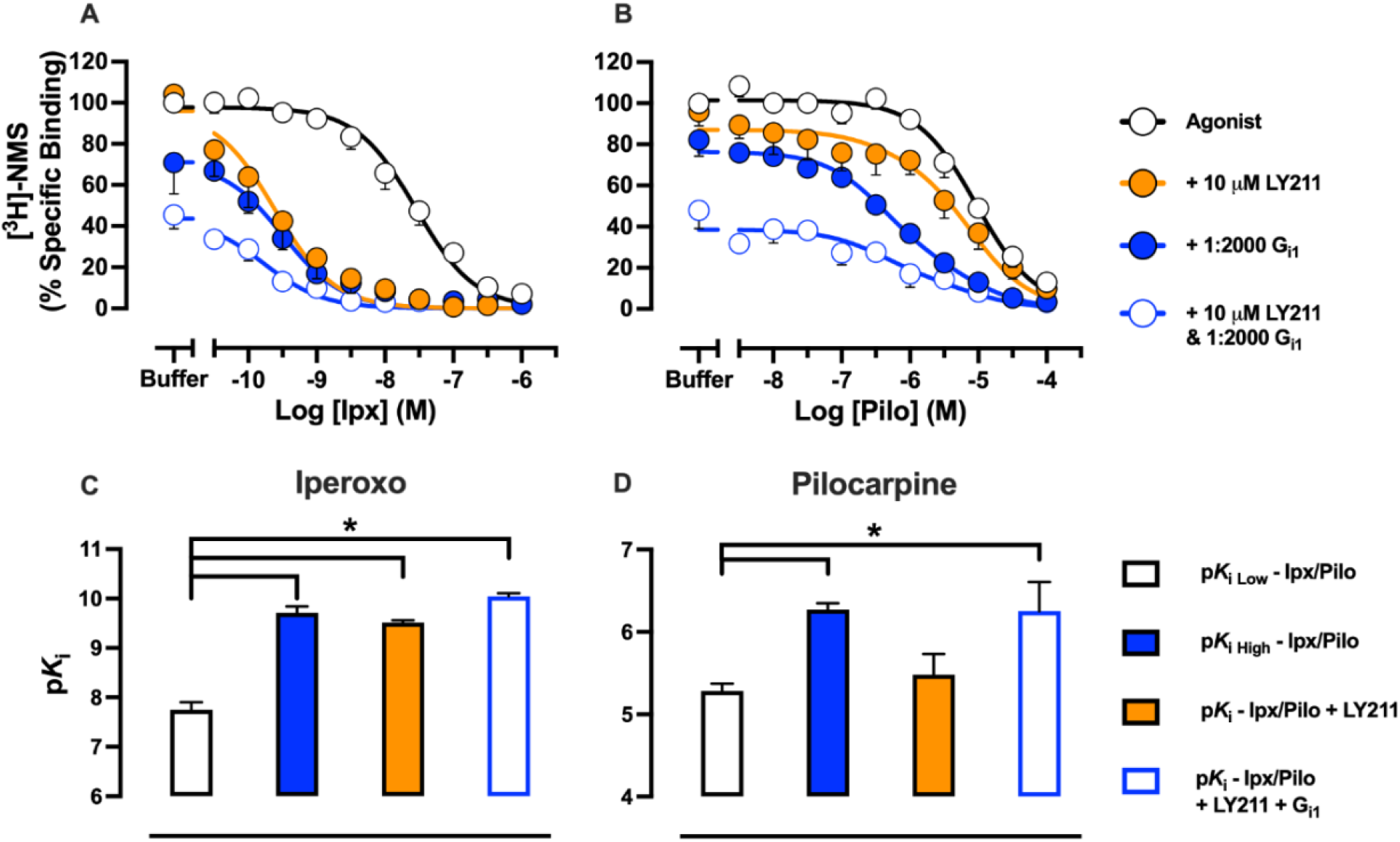
Modulation of the low and high-affinity state of super and partial agonists by G protein and PAM. Radioligand competition binding between **(A)** Ipx or **(B)** Pilo with [^3^H]-NMS at M_2_ mAChR nanodiscs in the presence of 10 μM LY2119620, a saturating stoichiometric 1:2000 ratio of receptor to G_i1_ proteins or 10 μM LY2119620 plus a saturating stoichiometric 1:2000 ratio of receptor to G_i1_ proteins. Data were normalised to the buffer only condition of Ipx or Pilo only and were fitted globally to a one-state model of competition binding. Low and high-affinity values of **(C)** Ipx obtained in **A** and **(D)** Pilo obtained in **B**. For all panels, data represent the mean ± S.E.M. of at least three individual experiments performed in duplicate. Group sizes and obtained parameters are listed in Table 1. *, significantly different, *p* < 0.05, one-way ANOVA, Tukey’s multiple comparisons test.

### Positive allosteric modulators kinetically stabilise high-affinity state

To further investigate the effect of PAMs on the high-affinity state, we turned to dissociation kinetic experiments to measure any kinetic influence the PAMs had on the high-affinity state. We pre-incubated the receptor with radioligand and added a saturating concentration of the orthosteric ligand atropine (Atr) to prevent rebinding of the radioligand, together with LY211 and/or G protein at different time points. Formation of the active state (through the addition of PAM or G protein) is expected to trap [^3^H]-NMS (*17, 41*), leading to a decrease in the dissociation rate of the radioligand. The extent of ternary complex stabilization by PAM or G protein will, therefore, be reflected in changes in [^3^H]-NMS dissociation rates. Without G protein, and in line with previous results (*40*), LY211 impeded the dissociation of the orthosteric radioligand (Fig. 5A-B, Table 1). Measuring dissociation in the presence of a saturating amount of G protein also led to a decrease in the dissociation rate of [^3^H]-NMS, consistent with a previous study that measured the impact of G_s_ on antagonist dissociation at the β_2_AR (*17*). Strikingly, combining both PAM and G protein produced a further decrease in the dissociation rate of the orthosteric radioligand (Fig. 5A-B, Table 1), suggestive of a synergistic closing of the orthosteric pocket where the PAM and G protein combine to facilitate the transition of receptors to the fully closed, active state. Given equilibrium affinity is a composite of kinetic association and dissociation rates and no changes in p*K*_i (High)_ at the M_2_ mAChR were seen with the addition of PAM, one would expect the association rate of an orthosteric ligand at the ternary complex to decrease in the presence of PAM. We pre-incubated the receptor with PAM and/or G protein and then added radioligand at different time points. Encouragingly, the G protein or PAM alone caused a decrease in the observed association rate of [^3^H]-NMS. The combination of both led to a large collapse and slowing down of the binding of [^3^H]-NMS, such that over the timescales employed, a large proportion of receptors were precluded from radioligand binding (as evidenced by a decrease in Bmax) (Fig. 5C-D, Table 1). Altogether, these findings suggest a synergistic kinetic stabilization of the active ternary complex by PAM and G protein.

**Figure 5:**
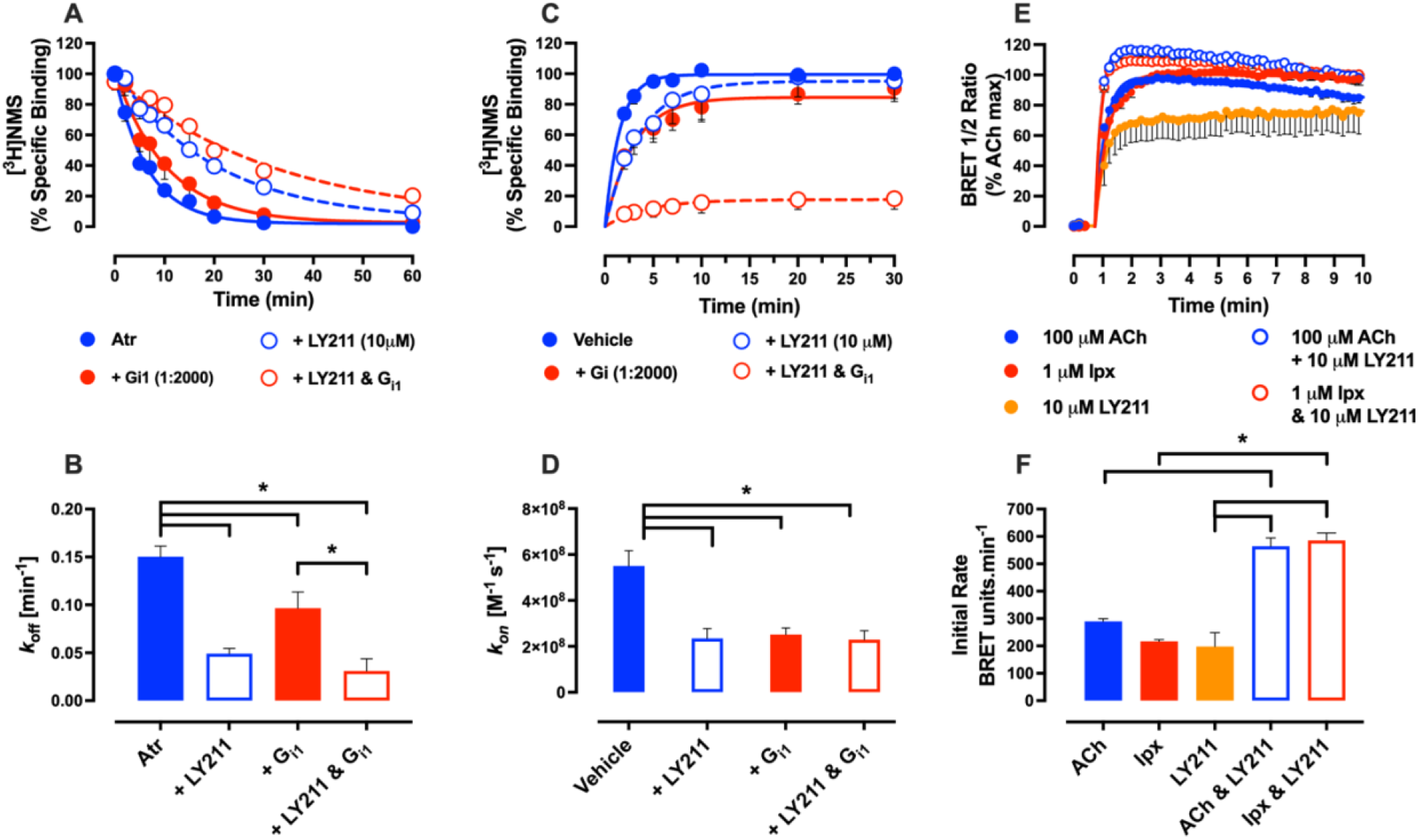
PAMs kinetically stabilise the high-affinity state of the ternary complex. **(A)** [^3^H]-NMS dissociation at M_2_ mAChR nanodiscs in the presence of atropine, atropine + 10 μM LY2119620, atropine + a saturating amount of G_i1_ and atropine + 10 μM LY2119620 + a saturating amount of G_i1_. Data were normalised to time point zero and were fitted globally to a one-phase exponential decay dissociation model. **(B)** Dissociation rates (*k*_off_ min^-1^) of data presented in A. **(C)** [^3^H]-NMS association at M_2_ mAChR nanodiscs in the presence of atropine, atropine + 10 μM LY2119620, atropine + a saturating amount of G_i1_ and atropine + 10 μM LY2119620 + a saturating amount of G_i1_. Data were normalised to time point zero and were fitted globally to a one-phase association model and constrained to *k*_off_ values obtained in A. **(D)** Association rates (*k*_on_ M^-1^ s^-1^) of data presented in C. **(E)** Kinetic G_i1_ protein activation at M_2_ mAChR FlpIn CHO cells transiently transfected with TRUPATH sensors in response to saturating concentrations of orthosteric and allosteric ligands. Data was baseline corrected to a buffer-only condition and normalised to ACh only, and a “Baseline then rise to steady state with drift” model was fit to the data. **(F)** Initial rate values, *v*_*o*_, of data presented in E. For panels A-F, data represent the mean ± S.E.M. of at least three individual experiments performed in duplicate. Group sizes and obtained parameters are listed in Table 1. *, significantly different, *p* < 0.05, one-way ANOVA, Tukey’s multiple comparisons test.

### Ternary complex stabilization leads to an increased rate of initial signalling

To determine whether the kinetic impact of the PAM on the ternary complex influences G protein activation, we turned to measuring the kinetics of G protein activation using the BRET TRUPATH assay in M_2_ mAChR FlpIn CHO cells. Using a recently described GPCR kinetic signalling model (*42, 43*), we derived an initial rate (*v*_*o*_) of signalling for saturating concentrations of ACh, Ipx, LY211 alone, and ACh or Ipx in combination with LY211. The term *v*_*o*_ is commonly utilised in enzyme kinetics and measures the rate of product formation prior to substrate degradation or establishment of equilibrium (the slope of the straight line of an activation curve). For a GPCR in a TRUPATH assay, this represents the efficiency of the agonist(s) occupied receptor at promoting G protein activation in the absence of any regulation or desensitisation mechanisms and can, therefore, be considered a direct reflection of the extent of active state stabilisation by the ligand(s).

ACh and Ipx displayed similar G protein activation kinetics (Fig. 5E-F, Table 1). Conversely, LY211 displayed slower G protein activation kinetics. However, when combining ACh and Ipx with LY211, a more rapid G protein activation was observed. A similar trend was observed in steady state values (which represent the final effect level stimulated by the ligand(s)), as ACh and Ipx displayed similar values that were both enhanced by LY211 (Fig. 5E-F, Table 1). Altogether, this indicates that the increased stabilisation of the ternary complex in the presence of orthosteric agonist and PAM leads to more efficient and rapid G protein activation.

## Discussion

GPCRs are inherently allosteric proteins owing to the distal communication that is required between the binding of an extracellular ligand and the activation of an intracellular transducer partner. Previous studies have shown that the allosteric coupling between G protein and orthosteric agonists leads to the formation of the ternary complex and high-affinity state that is structurally and pharmacologically characterised by the closing of the orthosteric pocket and increased orthosteric agonist affinity, respectively (*17, 24*). PAMs bind to a secondary ligand binding site, distinct from the endogenous ligand site, through which an allosteric transition enhances orthosteric agonist affinity. How a PAM influences the ternary complex and high-affinity state and the impact this has on GPCR signaling has remained largely unknown.

Through radioligand binding experiments with purified M_2_ mAChR nanodiscs and purified heterotrimer G protein, we show that once the high-affinity state is formed, the PAM LY211 does not promote a further increase in agonist affinity. This indicates that irrespective of orthosteric agonist efficacy, the agonist and G protein alone form the high-affinity state. The observation that a G protein is a superior PAM of agonist affinity compared to an exogenous PAM (LY211) may reflect the requirement of an allosteric link between the orthosteric binding site and G protein for GPCR activation and signalling. Our radioligand kinetic experiments show that the PAM LY211 stabilizes the lifetime of the ternary complex. Nanodiscs provide a suitable platform for the interrogation of the ternary complex due to the ability to control the presence of nucleotides and the concentration of each component. However, in a physiological setting, the ternary complex is transient due to the presence of freely available nucleotide. Measuring the kinetics of G protein activation in a recombinant cell environment and deriving a *v*_*o*_ parameter that represents the efficiency of the agonist-bound receptor (i.e., active state receptor), signalling parameters that capture agonist activity at the transient ternary complex were determined. These results show that PAMs promote the ability of orthosteric agonists to activate the G protein and, together with the observations from the nanodisc radioligand dissociation kinetics, indicate this arises from kinetic stabilisation of the ternary complex that leads to more efficient G protein binding and activation (Fig. 5E,F).

The discovery and development of allosteric modulators require the appropriate analysis and pharmacological quantification of parameters that capture the influence of allosteric modulators on orthosteric ligands and GPCR signalling. Through thermodynamic models, parameters reflective of equilibrium can be captured, including the ATCM that quantifies affinity (p*K*_B_) and binding cooperativity (log α) (*7, 39*) and the operational model of allosterism that quantifies functional cooperativity (log αb) and allosteric agonism (log ***τ***) (*32*). A kinetic ATCM has been developed to account for non-equilibrium artifacts in binding experiments due to the ability of allosteric modulators to slow the ligand-receptor interaction (*44, 45*). There is a growing appreciation for the kinetic aspects of ligand binding and GPCR signalling (*46*–*49*) with numerous studies linking ligand kinetics (*50, 51*) of orthosteric agonists to increased signalling (*52, 53*), biased agonism (*54, 55*), and the dynamics of conformational changes (*56*). However, the kinetic influence of allosteric modulators on GPCR signalling has remained relatively unexplored. Here, we have begun to explore this concept by utilizing a recently developed GPCR signalling kinetics model to show that PAMs enhance the initial rate of signalling at an orthosteric agonist-occupied receptor. Ultimately, this knowledge can be used to further explore the influence of allosteric modulators on GPCR signalling kinetics.

In the case of the adenosine A_1_ receptor (A_1_AR) reconstituted into nanodiscs, the PAM MIPS521 also slowed the dissociation rate of radioligand in the presence of G protein representative of ternary complex stabilization (*41*). Comparatively, the allosteric binding site of LY211 sits directly above and caps the orthosteric binding site, while the binding site of MIPS521 at the A_1_AR is distal to the orthosteric binding site sitting outside of the receptor at the lipid interface (Fig S7). These findings suggest a similar mechanism for stabilising the high-affinity state across Class A GPCRs irrespective of the position of the allosteric binding site. Differences exist between MIPS521 and LY211 in their ability to impede radioligand dissociation. For example, MIPS521 only impedes radioligand dissociation at A_1_AR nanodiscs in the presence of G protein, whereas LY211 can retard radioligand dissociation in both the presence and absence of G proteins. This can be rationalised by the differences in their binding site loci as the LY211 allosteric binding site is located directly above the orthosteric binding site, which sterically blocks orthosteric ligand dissociation. MIPS521, on the other hand, impedes dissociation solely by stabilising the high-affinity state as its allosteric binding site is distal to the orthosteric binding site.

One limitation of this study is that the ability of allosteric modulators to influence orthosteric ligand binding was explored experimentally through radioligand binding and conceptually through an ATCM. Within this framework, there is only one inactive state (low-affinity binding) and one active state (high-affinity binding) conformation. Though the majority of known allosteric modulators at the mAChRs operate through a two-state model of allosterism (*22*), questions remain over different forms of allosteric modulation. One such example is biased modulation, where allosteric modulators change the relative signalling capacity of an orthosteric agonist in one pathway over others (*57*). Numerous examples of biased modulation exist, and it is thought to occur through the presence of multiple agonist-bound active states, each linked to a different functional output, where the allosteric modulator changes the relative abundance of these active states (*58*). Indeed, at the M_2_ mAChR, analysing conformational dynamics through the use of ^13^CH_3_-ε-methionine nuclear magnetic resonance (NMR) it was shown that the M_2_ mAChR is highly dynamic and exhibits a wide range of receptor conformations (*59*) that are altered in the presence of LY211 (*60*). Similarly, at class C GPCRs, the metabotropic glutamate 2 (mGlu2) receptor exists in a number of conformational states when analysed through single-molecule Förster resonance energy transfer (smFRET) using fluorophores linked to SNAP tags in a detergent micelle environment (*61*). The presence of an agonist increased the fraction of receptors in the active state, and consistent with the results presented here, this approach found that the addition of a PAM led to increased stabilisation of the active state. Interestingly, this study also showed that the combined effect of PAM and G protein was not additive (*61*). A follow-up study focusing on the venus flytrap (VFT) domain indicated that PAMs stabilise the closed, active state of the VFT that is induced by agonists, (*62*) while a similar smFRET approach showed that the dynamics of the 7TM and cysteine-rich domain (CRD) were stabilized when the mGlu2 is co-bound to both agonist and PAM (*63*). We verify these findings at Class A GPCRs using radioligand binding experiments with receptors purified into a more native nanodisc environment. Furthermore, we extend this knowledge by linking active state stabilization of PAMs to increases in the signalling efficiency of the ternary complex in a recombinant cell environment. By tying together the thermodynamic and kinetic impacts of PAM and G protein on the stabilization of the active receptor state through novel biochemical and analytical approaches, our results provide further evidence of the molecular mechanisms that govern positive allosteric modulation at GPCRs.

## Materials and Methods

### Materials

Dulbecco’s modified Eagle medium (DMEM), Chinese hamster ovary (CHO) FlpIn cells were purchased from Invitrogen, Fetal bovine serum (FBS) was purchased from Thermotrace (Melbourne, Australia), hygromycin B was purchased from Roche Applied Science. [^3^H]-N-methylscopolamine ([^3^H]-NMS; specific activity, 70 Ci/mmol), MicroScint-O were purchased from PerkinElmer Life and Analytical Sciences (Waltham, MA, USA). The AlphaScreen-based *Sure-Fire*™ cellular extracellular signal-regulated kinase (ERK) 1/2 assay kit was purchased from TGR BioSciences (Adelaide, Australia). Prolume Purple was purchased from Nanolight Technologies (Pinetop, AZ, USA). All other chemicals were purchased from Sigma Chemical Company (St Louis, MO, USA).

### Mammalian Tissue Culture

FlpIn CHO cells stably expressing muscarinic acetylcholine receptor (mAChR) constructs were cultured at 37°C in 5% CO_2_ using DMEM supplemented with 5% (v/v) FBS. Upon reaching confluence, media was removed, cells were washed with phosphate buffered saline (PBS) and harvested from tissue culture flasks using versene. Cells were then pelleted through centrifugation at 350*g* for three minutes followed by resuspension in DMEM + 5% FBS. Cells were then either plated for an assay or reseeded into a tissue culture flask.

### ERK1/2 phosphorylation assay

The AlphaScreen-based SureFire kit was used for the measurement of phosphorylated ERK1/2 (pERK1/2). M_2ΔICL3_ mAChR FlpIn CHO cells were plated into 96-well plates at a density of 25 000 cells/well and incubated overnight at 37°C. The following day, the growth medium was replaced with serum-free DMEM medium for a minimum of 6h at 37°C. The cells were stimulated with a range of concentrations of drugs for an incubation period that matched the peak response time of the drug. Peak response time was determined by measuring pERK1/2 response at a range of time points over a 30 minute period. 10% FBS (v/v) was used as a positive control. Media was flicked off and cells were lysed with 80 μL/well SureFire lysis buffer and stored at -20°C overnight. Plates were thawed at room temperature (RT) and 10 μL of the cell lysates were transferred to a 384-well Optiplate. In reduced lighting conditions, 8.5 μl of detection buffer (buffer consisting of reaction buffer/activation buffer/acceptor beads/donor beads; in a 60:10:0.3:0.3 ratio) was added and plates were incubated for 1 hour at 37°C. Fluorescence signal was measured using an EnVision multilabel plate reader (PerkinElmer) with AlphaScreen settings. Data were expressed as a percentage of the pERK1/2 mediated by 10% FBS.

### G protein activation assay

Upon 60-80% confluence, wild type (WT) hM_2_ mAChR expressing FlpIn CHO cells were transfected transiently using Polyethylenimine (PEI, Sigma-Aldrich) with 10 ng per well of each the G protein TRUPATH biosensors pcDNA5/FRT/TO-Gα_i1_-RLuc8/pcDNA3.1-β_3_**/**pcDNA3.1-Gγ_9_-GFP2 giving a ratio of 1:1:1 ratio with 30 ng in total. Cells were plated at 30,000 cells per well into 96-well Greiner CELLSTAR white-walled plates (Sigma Aldrich). 48 hours later, cells were washed with 200 μL PBS and replaced with 1x HBSS supplemented with 10 mM HEPES. Cells were incubated for 30 minutes at 25C° before addition of 10 μL of 1.3 μM Prolume Purple coelenterazine (Nanolight technology, Pinetop, AZ). Cells were further incubated for 10 minutes at 25C° before bioluminescence resonance energy transfer (BRET) measurements were performed on a PHERAstar FSX plate reader (BMG Labtech) using 410/80-nm/515/30-nm filters. Four baseline measurements for each well were taken before addition of drugs or vehicle to give a final assay volume of 100 μL and ligand induced changes in BRET measurements were then taken for and additional 10 minutes. BRET signal was calculated as the ratio of 515/30-nm emission over 410/80-nm emission. The ratio was vehicle corrected using the initial four baseline measurements and then baseline corrected again using the vehicle-treated wells. Concentration response curves were constructed from the area under the curve (AUC) of the double baseline corrected kinetic traces and normalised to the highest AUC value in each dataset. For kinetic TRUPATH experiments, the double baseline corrected BRET ratios were normalised to the greatest ACh induced BRET decrease in each dataset.

### Membrane Preparation

At confluence, WT hM_2_ mAChR FlpIn CHO cells were harvested and pelleted. The pellet was resuspended in HEPES homogenisation buffer (50 mM HEPES, 2.5 mM MgCl_2_ and 2 mM EGTA) and homogenized for three 10 second intervals with a 30 second cooling interval on ice. The homogenate was centrifuged for 10 minutes at 600*g* and the supernatant was stored on ice. The remaining pellet was resuspended in HEPES homogenisation buffer and the homogenization process was repeated until no further reduction in pellet size occurred following centrifugation. The supernatant was placed into centrifugation tubes and centrifuged (30,000*g*, 30 minutes, 4°C) and the pellet was resuspended into binding buffer (20 mM HEPES, 100 mM NaCl, 10 mM MgCl_2_). Protein concentration was determined using the Pierce BCA protein assay kit, ThermoFisher.

### ApoA1 cMSP1D1 expression and purification

E. coli BL21 (DE3) cells were transformed with split intein cMSP1D1 constructs inserted into pET28a vector (Novagen) in LB media + kanamycin. Induction of protein expression was accomplished through 500 mM Isopropyl β-d-1-thiogalactopyranoside (IPTG) at an OD600 of 0.6 and cells were shaken for 16 to 20 hours at 25°C. Cells were harvested by centrifugation (7000*g*, 20 minutes, 4°C) and the cell pellet was resuspended in lysis buffer (50 mM Tris pH 8.0, 250 mM NaCl, 0.5% Triton X-100), 0.5 mM EDTA, 1 mM phenylmethylsulfonyl fluoride (PMSF) were added and the cells were lysed by incubation with 50 μg/mL lysozyme for 30 minutes and further sonication. Lysate was dounced and put though Avestin followed by a 30 minute incubation at 4 °C with 2 μL benzonase + 10 μM benzamidine and 5 mM MgCl_2_. Cell debris were removed by centrifugation (30 000*g*, 30 minutes, 4°C). A heat shock at 70°C was conducted with the soluble fraction for 40 minutes. Aggregates were removed by centrifugation (30 000*g*, 30 minutes, 4°C). The supernatant was loaded onto a gravity flow Ni-NTA resin column (GE healthcare). Flow-through and 1 column volume (CV) of wash buffer (20 mM Tris pH 8.0, 320 mM NaCl, 10 mM imidazole, 10 mM 2-Mercaptoethanol (BME)) were collected and dialyzed to 20 mM Tris pH 8.0, 0.5 mM EDTA, 10 mM BME. The urea concentration was set to 6 M and the sample was applied to a 5 mL HiTrap QFF anion exchange column (GE healthcare) and was eluted using a 30 CV long gradient from low salt buffer (20 mM Tris pH 8.0, 0.5 mM EDTA, 6 M urea, 10 mM BME) to high salt buffer (20 mM Tris pH 8.0, 300 mM NaCl, 0.5 mM EDTA, 6 M urea, 10 mM BME). Pure protein was pooled, dialysed to 20 mM Tris pH 8.0, 200 mM NaCl, 0.5 mM EDTA, 10 mM BME, concentrated using a 10 kDa molecular mass cut-off centrifugal filter unit (Millipore, Burlington, MA, USA), flash frozen using liquid nitrogen and stored at -80°C.

### M_2_ mAChR expression and purification

The human M_2_ muscarinic receptor gene (http://www.cdna.org) was modified to give a receptor containing an N-terminal anti-Flag epitope tag and a carboxy (C)-terminal 8× histidine tag. In order to increase stability and expression, residues 226–389 of ICL3 were removed. The resulting Flag-M_2ΔICL3_-His construct was cloned into a PVL1392 baculovirus transfer vector. M_2_ mAChR protein were expressed using the Bac-to-Bac Baculovirus Expression System (Invitrogen) in Sf9 cells. Cells were grown in ESF 921 serum-free media (Expression System) and infected at a density of 4.0 × 10^6^ cells per millilitre, treated with 10 μM atropine (Atr) and shaken at 27°C for 48-60 hours. Cells were harvested by centrifugation (10 000*g*, 20 minutes, 4°C) and cell pellet resuspended in lysis buffer (10 mM Tris, pH 7.5, 1 mM EDTA, 1 mM MgCl_2_, protease inhibitors (500 μM PMSF, 1 mM LT, 1 mM benzamidine), 1 mg/mL iodoacetamide, benzonase and 1 μM Atr) and stirred at 25°C until homogenous. Cell lysate was centrifuged (15 000 rpm, 15 minutes, 4°C). Receptor was solubilized in solubilisation buffer (30 mM HEPES pH 7.5, 1% n-Dodecyl-B-D-Maltoside (DDM), 0.2% cholate, 0.03% cholesterol hemisuccinate (CHS), 750 mM NaCl, 30% glycerol, protease inhibitors (500 μM PMSF, 1 mM LT, 1 mM benzamidine), 1 mg/mL iodoacetamide, benzonase and 1 μM Atr). Soluble fraction was separated through centrifugation (15 000 rpm, 15 minutes, 4°C) and supernatant was incubated with Ni-NTA resin for 2h at 4°C. Ni-NTA resin was washed with wash buffer (30 mM HEPES pH 7.5, 0.1% DDM, 0.02% sodium cholate, 0.003% CHS, 750 mM NaCl, 30% glycerol, 5 mM imidazole and 1 μM Atr) and protein was eluted with wash buffer supplemented with 250 mM imidazole. Sample was loaded onto M1 anti-Flag affinity resin and detergent was exchanged from DDM solubilisation buffer to Lauryl maltose-neopentyl glycol (LMNG) buffer (30 mM HEPES pH 7.5, 0.01% LMNG, 0.001% CHS, 100 mM NaCl and 1 μM NaCl. Protein was eluted of M1 anti-Flag affinity resin through LMNG buffer supplemented with 10 mM EDTA and 0.2 mg/mL Flag peptide. Elution was concentrated and run through size exclusion chromatography (SEC) using Superdex200 increase 10/300 column (GE Healthcare) with LMNG buffer. Sample was collected, concentrated and flash frozen using liquid and stored at -80°C.

### G Protein expression and purification

WT G protein alpha subunits were cloned into a PVL1392 baculovirus transfer vector. G protein beta and gamma subunits were cloned into a PVL1392 baculovirus transfer vector with the beta subunit modified to contain a carboxy (C)-terminal 8× histidine tag. G protein subunits were expressed using the Bac-to-Bac Baculovirus Expression System (Invitrogen) in Trichoplusia ni (Hi5) insect cells. Cells were grown in ESF 921 serum-free media (Expression System) and infected at a density of 4.0 × 10^6^ cells per millilitre with a 1:1 ratio of Gα to Gβγ viruses and shaken at 27°C for 48-60 hours. Cells were harvested by centrifugation (10 000*g*, 20 minutes, 4°C) and cell pellet was lysed in lysis buffer (10 mM Tris pH 7.4, 5 mM MgCl_2_, 5 mM tris(2-carboxyethyl)phosphine (TCEP), and 10 μM guanosine 5’-diphosphate (GDP), + protease inhibitors (500 μM PMSF, 1 mM Leupeptin, 1 mM benzamidine), benzonase). Cell lysate was centrifuged (18 000 rpm, 15 minutes, 4°C). Cell pellet was solubilised in 20 mM HEPES pH 7.5, 100 mM NaCl, 1.0% sodium cholate, 0.05% DDM, 5 mM MgCl_2_, 1 mM TCEP, 10 μM GDP, protease inhibitors (500 μM PMSF, 1 mM LT, 1 mM benzamidine), 20 mM imidazole and stirred for 60 minutes at 4°C followed by centrifugation (18 000 rpm, 15 minutes, 4°C). Filtered supernatant was incubated with Ni-NTA resin for 90 minutes at 4°C. Resin was loaded onto glass column and washed with wash buffer (20 mM HEPES pH 7.5, 100 mM NaCl, 0.05% DDM, 1 mM MgCl_2_, 1 mM TCEP, 10 μM GDP, protease inhibitors (500 μM PMSF, 1 mM LT, 1 mM benzamidine, 20 mM imidazole), until no more protein was coming off as determined by Bradford.

Sample was eluted with wash buffer + 250 mM imidazole. Sample was dialysed overnight at 4°C to remove imidazole and to lower NaCl to 150 mM. The following morning the sample was loaded onto 5 mL HiTrap QFF anion exchange column (GE healthcare) and then washed with 15 CV of buffer A (20 mM HEPES pH 7.4, 25 mM NaCl, 0.1% DDM, 1 mM MgCl_2_, 100 μM TCEP, 10 μM GDP). A gradient of 0-30% over 20 CV was then run with buffer A and buffer B (20 mM HEPES pH 7.4, 1 M NaCl, 0.1% DDM, 1 mM MgCl_2_, 100 μM TCEP, 10 μM GDP). Samples were collected, diluted with 20 mM HEPES pH 7.4, 30 mM NaCl, 0.1% DDM, 1 mM MgCl_2_, 100 μM TCEP, 10 μM GDP to dilute NaCl to a final concentration of 125 mM. Sample was concentrated, glycerol added to 20%, flash frozen using liquid nitrogen and stored at -80°C.

### Incorporating M_2_ mAChR into nanodiscs

1-palmitoyl-2-oleoyl-sn-glycero-3-phosphocholine (POPC, Avanti Polar Lipids) and 1-palmitoyl-2-oleoyl-sn-glycero-3-phospho-(1’-rac-glycerol) (POPG, Avanti Polar Lipids) were mixed using chloroform to a final concentration of 10 mM POPC and 6.67 mM POPG (3:2 POPC:POPG ratio). The lipid mixture was dried using nitrogen to evaporate the chloroform, residual moisture and any remaining chloroform was then removed through the use of an overnight incubation step in a desiccator. The following morning, the lipid mixture was resuspended in HNE buffer (20 mM HEPES pH 8.0, 100 mM NaCl, 1 mM EDTA, 50 mM sodium cholate) and sonicated on ice until the lipid solution became clear. Lipid mixture was flash frozen using liquid nitrogen and stored under nitrogen at -80°C. To determine optimum ratio of lipid to ApoA1, different ratios of ApoA1 to lipid (1:40, 1:40, 1:60, 1:70 and 1:80) were tested with 100 μM ApoA1 to determine the optimum ratio that created a homogenous, monodisperse sample as determined by SEC using a Superdex200 increase 10/300 column (GE Healthcare). In brief, the appropriate amount of lipid mixture, HNE buffer (20 mM HEPES pH 8.0, 100 mM NaCl, 1 mM EDTA), 100 μM ApoA1 in a 100 μL volume were mixed together. Following incubation on ice for 1h, the mixture was incubated with ∼50 mg of Bio-Beads (Bio-Rad) to remove all detergents and initiate the spontaneous formation of nanodiscs. The sample was mixed overnight at 4°C and the following morning the nanodiscs were purified through use of size-exclusion chromatography on a Superdex200 increase 10/300 column (GE Healthcare) with HNE buffer. To incorporate purified receptor, the appropriate lipid ratio was chosen, and nanodisc reconstitution was performed as described above with the inclusion of 5 μM purified receptor. Samples were pooled, flash frozen using liquid nitrogen and stored at -80°C. For nanodiscs that were reconstituted with M_2_ mAChR purified with Atr, the sample was dialyzed for 24h at 4°C with PBS prior to flash freezing to remove residual Atr.

### Fab complexing

Pierce Fab Preparation Kit (ThermoFisher) was used to generate Fab fragments of anti-Flag IgG antibody. Purified Fab fragments were added to M_2_ mAChR cMSP1D1 at a ratio of 10:1, where the concentration of receptor in the nanodisc was calculated through saturation radioligand binding with [^3^H]-NMS (described in general methods). The sample was incubated with 5 mM MgCl_2_ and incubated at RT for 4 hours. The sample was loaded over 0.5 mL of Ni resin, flow-through collected and reloaded for a total of five times. The resin was washed until no more protein was coming off the column as determined by Bradford and the sample was eluted with PBS + 250 mM imidazole. Sample were concentrated analysed through negative staining.

### Radioligand binding

The affinity of [^3^H]-NMS for M_2_ mAChR in nanodiscs, as well as the concentration of receptor in nanodiscs, was determined through saturation binding with [^3^H]-NMS. M_2_ nanodiscs were incubated with a range of concentrations of [^3^H] NMS in a final volume of 200 μL binding buffer (20mM HEPES pH 7.5, 100mM NaCl, 10mM MgCl_2_, 0.5% BSA) for 6 hours at RT. To determine low and high-affinity of orthosteric agonists at M_2_ mAChR nanodiscs, nanodiscs were incubated in a final volume of 200 μL binding buffer containing a range of concentrations of the cold ligand in the presence of a K_D_ concentration of [^3^H]-NMS as determined through saturation binding in the presence of a saturating/increasing stoichiometric amounts of G protein for 6 hours at RT. Concentrated G protein was added such that DDM and GDP were diluted at least 1600-fold. For interaction experiments with LY211 competition binding between a K_D_ concentration of [^3^H]-NMS and a range of concentrations of an orthosteric drug was performed in the presence of varying concentrations of LY211 with or without a saturating amount of purified G protein. For dissociation experiments, M_2_ mAChR nanodiscs were incubated with a K_D_ concentration of [^3^H]-NMS for 1 hour at RT in a total volume of 200 μL of binding buffer. Dissociation of the radioligand was initiated by addition of 10 μM Atr alone or in the presence of LY211 and G protein at various timepoints. For association experiments, M_2_ mAChR nanodiscs were incubated with LY211 and G protein for 4 hours at RT before association was initiated through addition of a K_D_ concentration of [^3^H]-NMS at various timepoints. Membrane equilibrium binding experiments were performed with 15 μg of membrane expressing the WT hM_2_ mAChR in a 300 μL reaction volume of binding buffer with or without 100 μM 5’Guanylyl imidodiphosphate (Gpp(NH)p) in 96w deep well plates for 6 hours at RT. Membranes were incubated with a range of concentrations of ACh in the presence of a K_D_ concentration of [^3^H]-NMS as determined through saturation binding. For all radioligand binding experiments, non-specific binding was defined by 10 μM of Atr and the assay was terminated through rapid filtration through Whatman GF/B filter plates using a 96-well harvester (Perkin-Elmer). Filter plates were dried overnight and radioactivity was determined by the addition of 40 μL of MicroScint-O scintillation fluid and counting in a MicroBeta^2^ Plate Counter (PerkinElmer Life Sciences).

### Data Analysis

All data were analysed using GraphPad Prism 9 (Graphpad Software, San Diego, CA). Concentration response curves were fit with a three-parameter logistic equation to quantify potency (pEC_50_) and an operational model of agonism to derive an efficacy parameter (log ***τ***) (*64*). The interaction between ACh and LY211 in G protein activation assay was fit to an operational model of allosterism to derive functional modulation (log αβ) and affinity (p*K*_B_) parameters (*32*). For competition binding experiments between [^3^H]-NMS and a range of concentrations of unlabelled orthosteric agonist at M_2_ mAChR nanodiscs in the presence of G protein, and for competition binding at M_2_ mAChR membranes, data was fit to a two-site competition binding model or a one-site competition binding model. For the interaction of ACh with a range of LY211 concentrations with and without a saturating amount of G protein, data was fit to an allosteric ternary complex model to derive p*K*_B_, and α binding cooperativity parameters (*39*). Radioligand dissociation and association data at M_2_ mAChR nanodiscs was fit to mono-exponential decay and one-phase association equations, respectively. Kinetic TRUPATH data was analysed using a ‘baseline then rise to steady state with drift’ equation (*42*). All affinity, potency, cooperativity and efficacy parameters were estimated as logarithms, and statistical analysis between different treatment conditions were determined by one-way ANOVA using a Tukey’s multiple comparisons with a value of *p* < 0.05 considered as significant.

## Acknowledgments

We thank Sam R. J. Hoare for his discussion on use of the GPCR kinetic signalling model. Figures 1B and 3D,G,H were created with BioRender.com.

## Contributions

Conceptualization: WACB, CJDJ, AC, DMT. Methodology: WACB, CJDJ, CV, AC, DMT. Investigation: WACB. Funding acquisition: CV, AC, DMT. Supervision CV, AC, DMT. Writing – original draft: WACB, AC, DMT. Writing – review & editing: WACB, CJDJ, CV, AC, DMT.

## Data and materials availability

All data needed to evaluate the conclusions in the paper are present in the paper and/or the Supplementary Materials.

