## Supplemental Data for "Positive allosteric modulation of a GPCR ternary complex"

Wessel A. C. Burger *et al.*

**This PDF file includes:**

Figs. S1 to S7

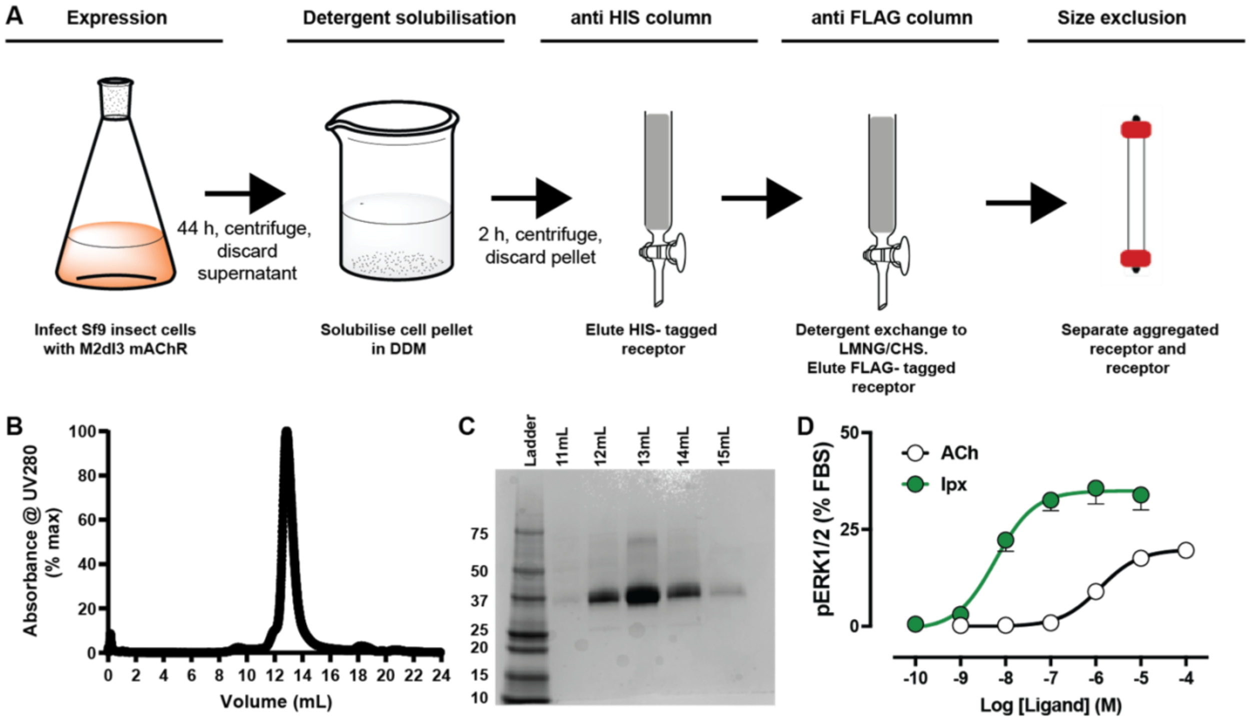


**Fig. S1. Expression and purification of $M_{2\Delta ICL3}$ mAChR. (A)** Expression and purification scheme of $M_{2\Delta ICL3}$ mAChR **(B)**. SEC trace of purified $M_{2\Delta ICL3}$ mAChR. **(C)** Coomassie staining of fractions obtained in **B**. **(D)** pERK phosphorylation concentration-response curves for ACh and Ipx at the $M_{2\Delta ICL3}$ mAChR expressed in FlpIn CHO cells. Data are normalized to FBS response and represent the mean ± s.e.m. of seven independent experiments performed in duplicate. Data are globally fitted to a three-parameter logistic equation to obtain $pEC_{50}$ values of ACh: 5.91 ± 0.09 and Ipx: 8.23 ± 0.16.

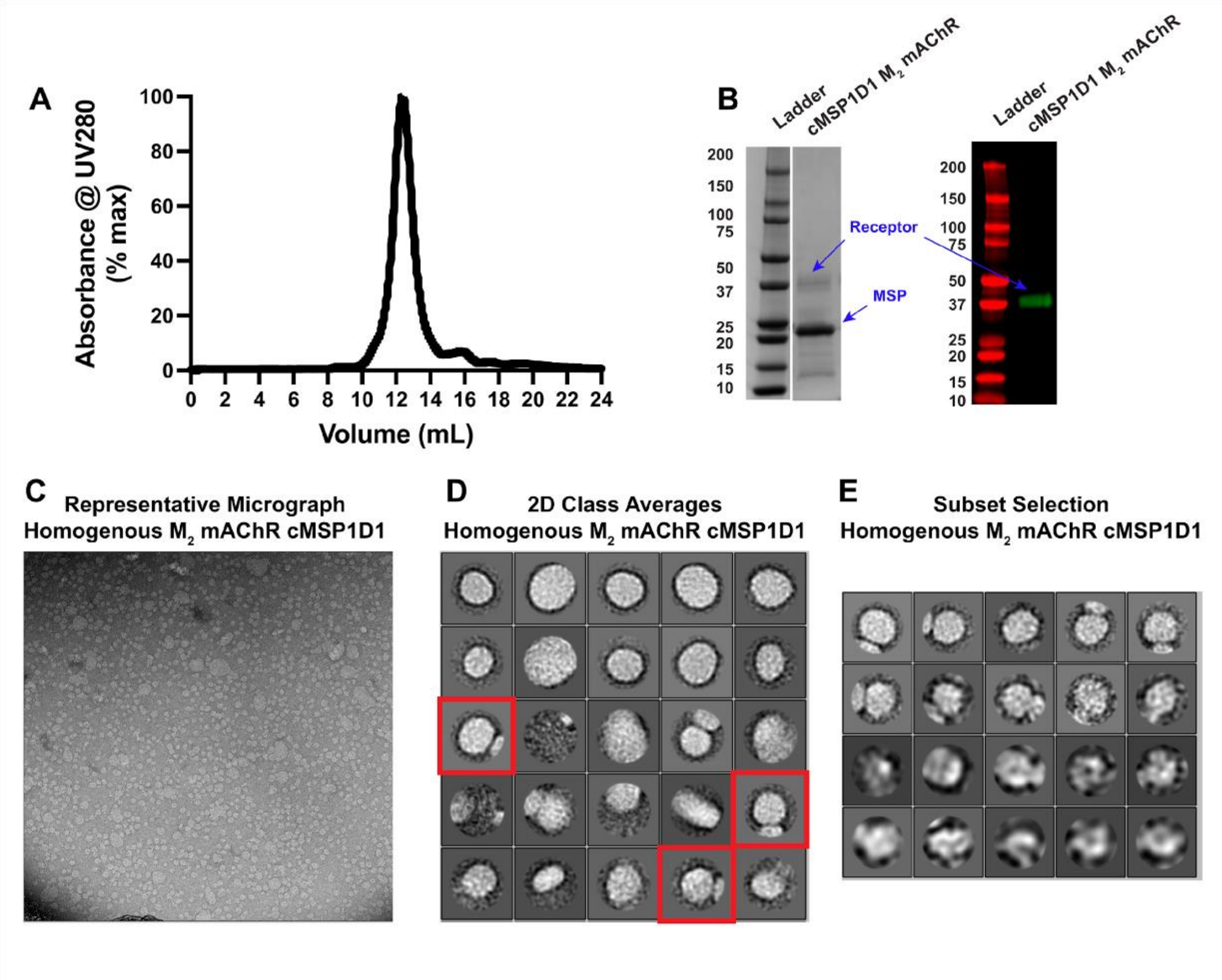


**Fig. S2. Reconstitution of $M_2$ mAChR into nanodiscs. A)** SEC traces following $M_2$ mAChR reconstitution into cMSP1D1 nanodiscs. **B)** The presence of receptor and MSP in each nanodisc was verified through left SDS-Page with Coomassie staining and right western blotting with anti-Flag antibody. **C)** Representative negative staining micrograph of M2 mAChR nanodiscs complexed with anti-Flag FAB **D)** 2D class averages for the $M_2$ mAChR+Fab cMSP1D1 sample. 2D classes surrounded by a red border were selected and used to perform further subset selection 2D classification in **E)**.

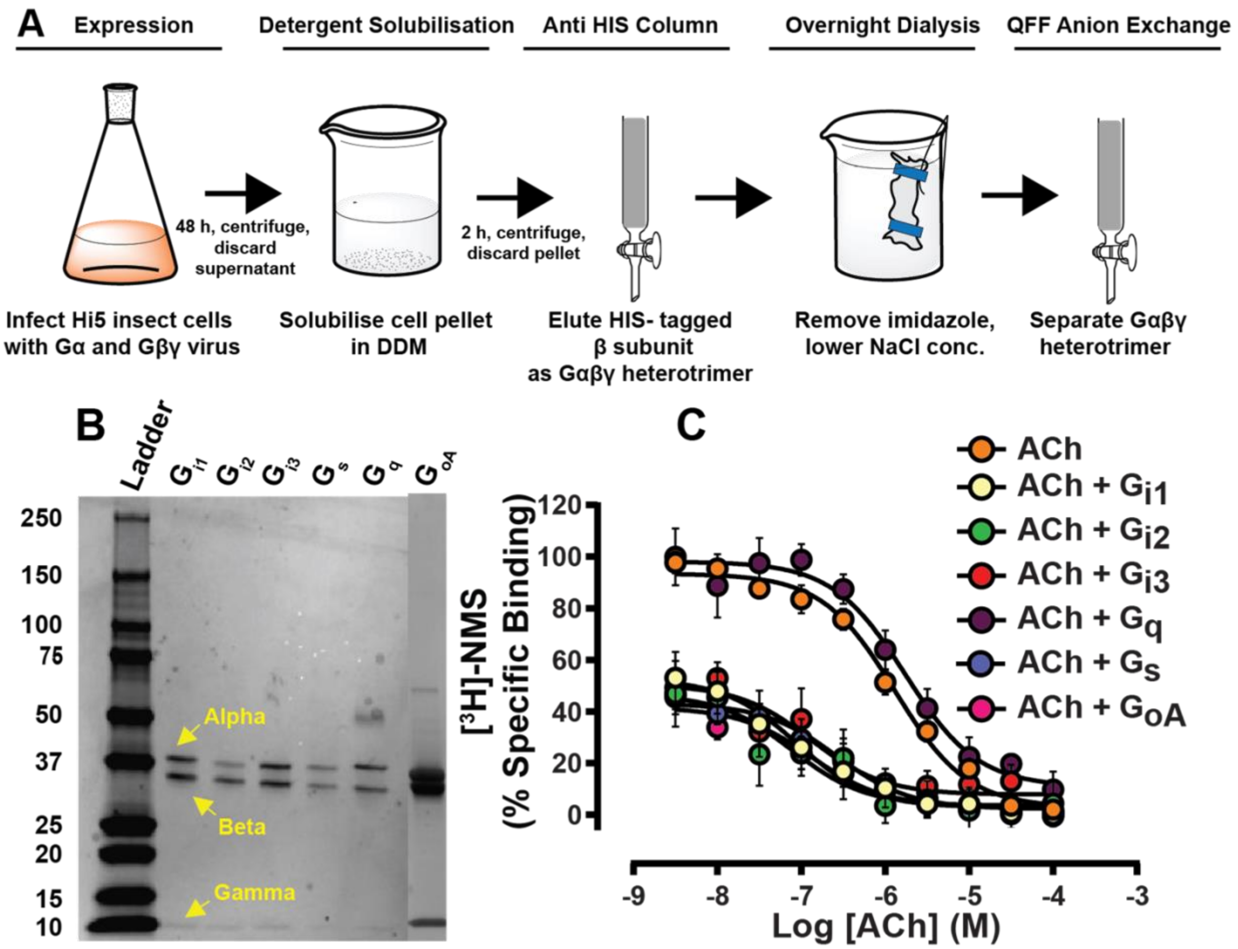


**Fig. S3. Expression, purification and characterisation of WT G protein heterotrimers. (A)** Expression and purification scheme for WT G protein heterotrimers. **(B)** Coomassie staining of purified WT G protein heterotrimers. **(C)** Competition binding of ACh and [$^3$H]-NMS in the presence of a saturating amount of G protein at $M_2$ mAChR cMSP1D1 nanodiscs. Data represent the mean ± s.e.m. of at least three independent experiments performed in duplicate.

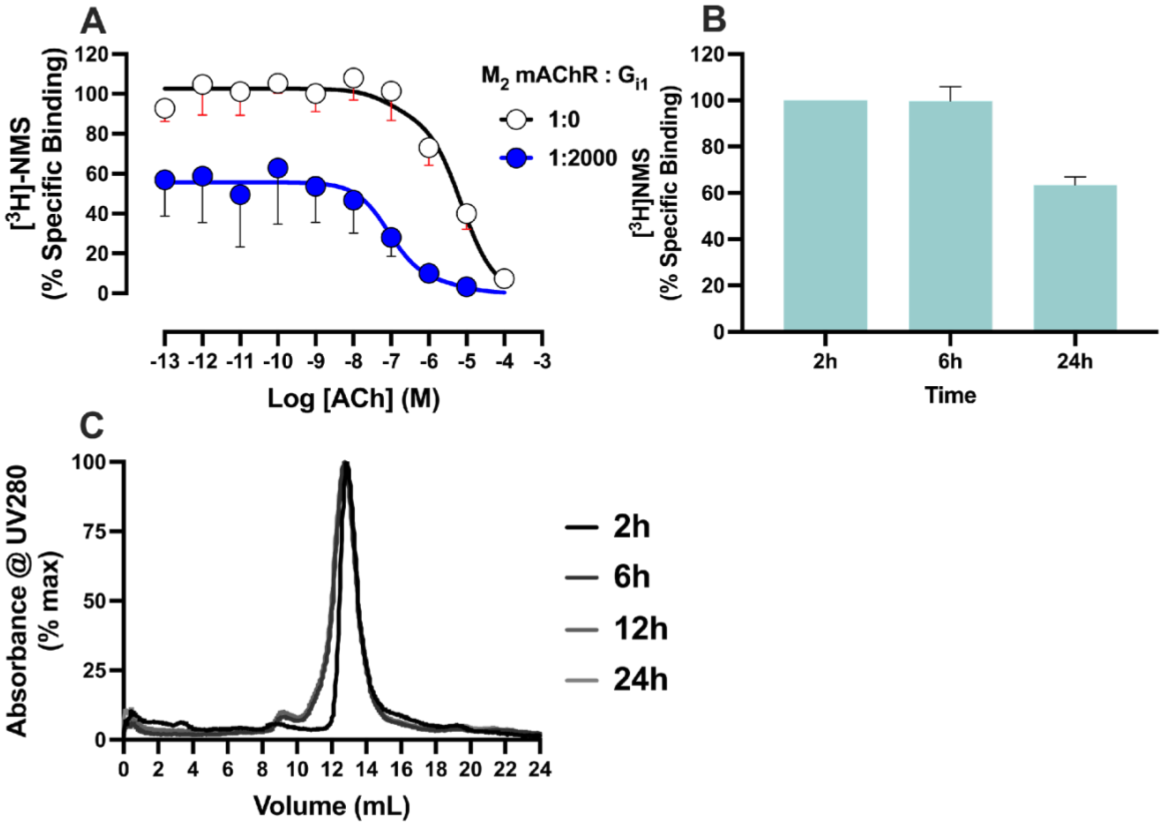


**Fig. S4. Stability of $M_2$ mAChR nanodiscs. (A)** Radioligand competition binding following 4 hours at room temperature between ACh and [$^3$H]-NMS at $M_2$ mAChR nanodiscs without and with a saturating amount $G_{i1}$ protein. Data were normalised to the buffer only condition and represent the mean ± S.E.M of three individual experiments performed in duplicate. Data are globally fitted to a two-state model of competition binding where p$K_{i\,(Low)}$ and p$K_{i\,(High)}$ values were shared. p$K_{i\,(Low)}$: 5.44 ± 0.31 and p$K_{i\,(High)}$: 7.37 ± 0.57 values for ACh were obtained. **(B)** Percentage of binding of a $K_D$ concentration of [$^3$H]-NMS over time at $M_2$ mAChR nanodiscs. Data is normalized to the level of [$^3$H]-NMS binding observed at 2 hours. **(C)** SEC traces $M_2$ mAChR nanodiscs following incubations of different time periods at room temperature.

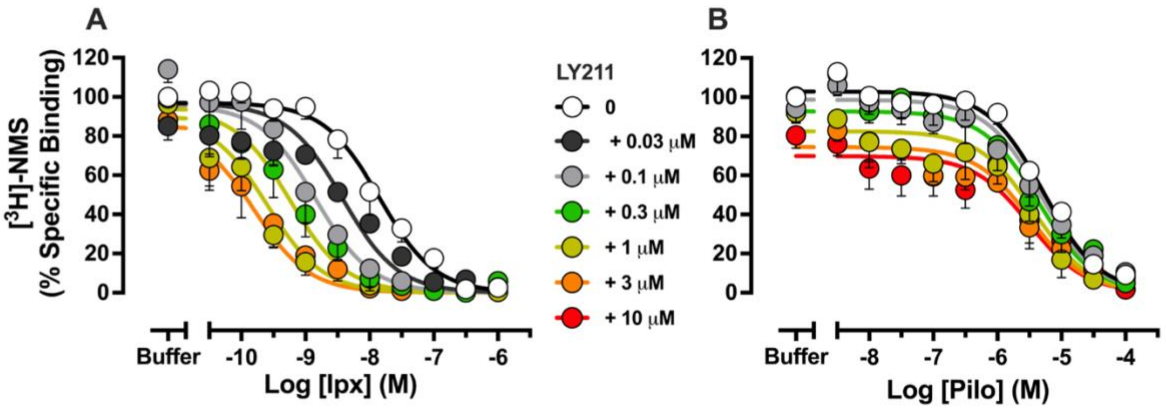


**Fig. S5. Interaction binding of iperoxo and pilocarpine with LY2119620.** Radioligand competition binding between **(A)** Ipx or **(B)** Pilo and [$^3$H]-NMS with increasing amounts of concentrations of LY211 at $M_2$ mAChR nanodiscs. For all panels, data represent the mean ± S.E.M. of at least three individual experiments performed in duplicate. Data were normalised to the buffer only condition and were fitted globally to an allosteric ternary complex model. Group sizes and obtained parameters are listed in Table 1.

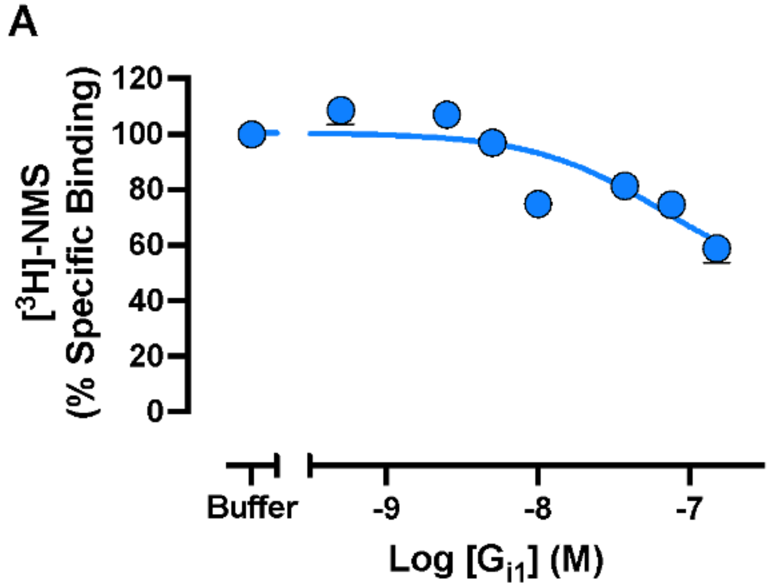


**Fig. S6. $G_{i1}$ heterotrimer protein versus [$^3$H]-NMS.** Binding of [$^3$H]-NMS at $M_2$ mAChR nanodiscs in the presence of increasing concentrations $G_{i1}$ heterotrimer. Data is replotted from the buffer only condition of Figures **2A**, **4A** and **4B** with G protein ratios converted to molar concentrations. Data were normalised to the buffer-only condition and were fitted globally to an allosteric ternary complex model. Group sizes and obtained parameters are listed in Table 1.

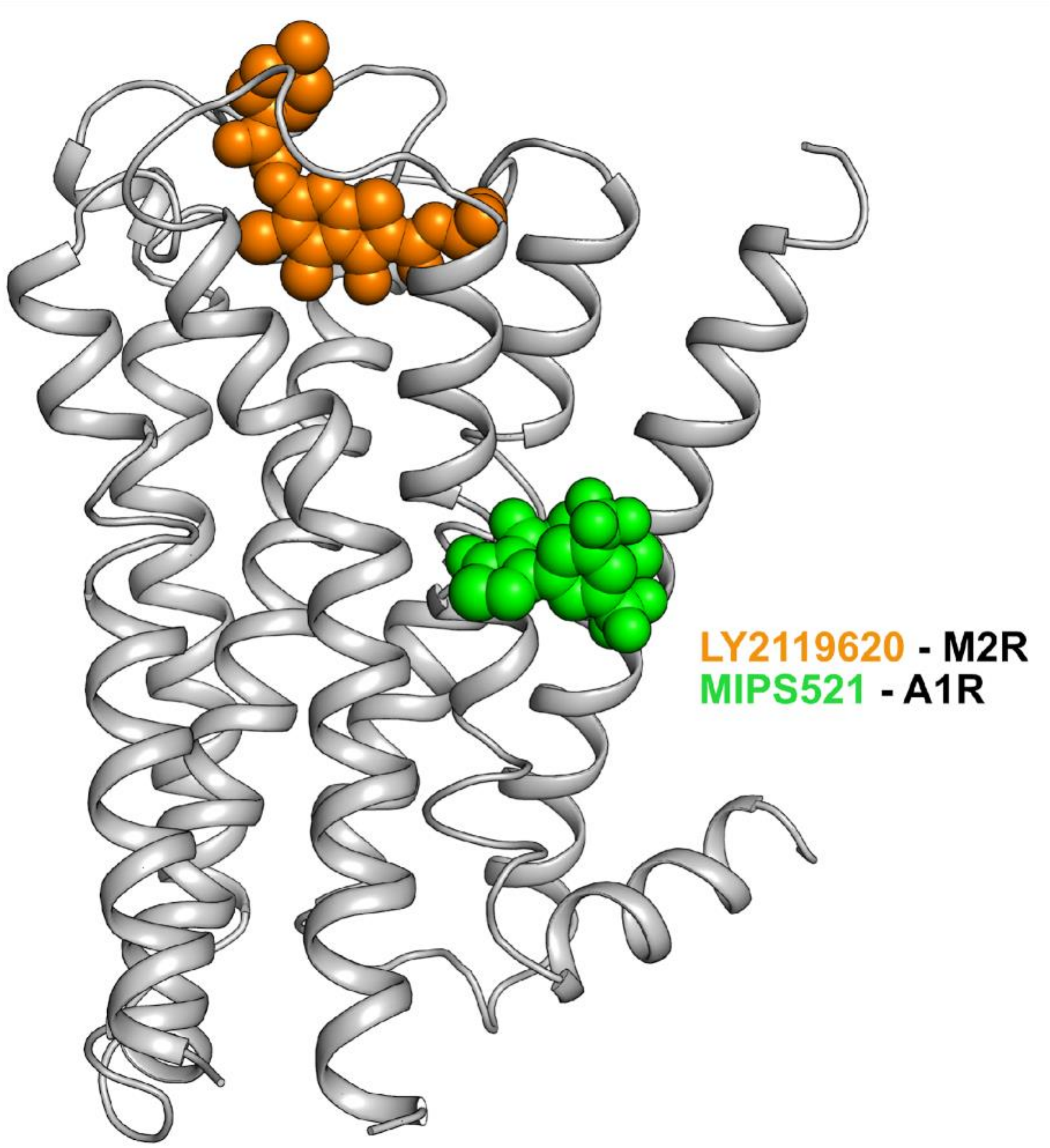


**Fig. S7. Location of LY2119620 and MIPS521 binding sites.** Comparison of the LY2119620 binding loci at the $M_2$ mAChR (PDB: 4MQT) and the MIPS521 binding loci at the A1AR (PDB: 7LD3).
